# Smooth muscle cell estrogen receptor alpha promotes arterial stiffness in the absence of estradiol

**DOI:** 10.64898/2026.03.03.709417

**Authors:** Casey G. Turner, Jacqueline Matz, Jade Breton, Karla C. de Oliveira, Rachel Kenney, Jennifer Vorn, Michelle Zhao, Jaime Ibarrola, Qing Lu, Gregory Martin, Zhe Sun, Iris Z. Jaffe, Michael A. Hill, Jennifer J. DuPont

## Abstract

**Background:** Clinical evidence supports a greater impact of arterial stiffening in cardiovascular mortality in women versus men. Arterial stiffness increases across the menopausal transition, implicating a role of the loss of estrogens in arterial stiffening, but mediating mechanisms remain unclear.

**Methods:** The role of estradiol and smooth muscle cell (SMC) estrogen receptor alpha (ERα) in arterial stiffening, by aortic pulse wave velocity (PWV), was assessed in 3 models: (1) the loss of estradiol in young, female mice comparing sham surgery or bilateral ovariectomy (OVEX) ± estradiol, (2) the impact of sham versus OVEX surgery in young, female SMC-ERα-intact and SMC-ERα-knockout (KO) littermates, and (3) arterial stiffening during natural aging by comparing young and aged, female and male SMC-ERα-intact and SMC-ERα-KO littermates. Mechanistic pathways were assessed using histological assessment of aortic fibrosis and elastin degradation, aortic MMP expression, and atomic force microscopy.

**Results:** OVEX increased PWV and aortic medial fibrosis, with no impact on elastin integrity, in young female mice. Arterial stiffening and fibrosis were prevented in OVEX mice that were supplemented with estradiol. OVEX-induced arterial stiffening in SMC-ERα-intact female mice was prevented in SMC-ERα-KO littermates. In this model, OVEX was also associated with increased aortic medial fibrosis without changes in elastin integrity. Aging from 3 to 18 months significantly increased PWV in female and male SMC-ERα-intact mice. Aging-induced stiffening was fully prevented in female and partially prevented in male SMC-ERα-KO mice. SMC-ERα contributes to aging-associated arterial stiffening by sex-specific mechanisms, including elastin degradation in females and phenotypic changes in SMC stiffness and probability to form cellular adhesions in males. Circulating estradiol was significantly decreased in serum from aged compared with young female mice.

**Conclusions:** These findings support that SMC-ERα contributes to arterial stiffening in female and male mice in situations where the vasculature is exposed to low levels of estradiol.

## Introduction

Arterial stiffening occurs with aging and is an independent predictor of hypertension incidence, major cardiovascular events, and mortality^1–3^. There are sex-dependent differences in arterial stiffness across the lifespan, such that arterial stiffness is lower in women compared with men until approximately the 5^th^ to 6^th^ decade of life, after which arterial stiffening occurs at a greater rate and reaches greater absolute values in elderly women compared with men^4^. Additionally, arterial stiffness has a ∼2-fold greater impact on all-cause and cardiovascular-specific mortality in women compared with men, which is exacerbated to over 3-fold when women are older than 55 years of age^5^. The dramatic changes in arterial stiffening and cardiovascular risk occur after an average age of menopause in women, which is between 51 and 52 years of age^6^. Clinical observations further support that the rate of arterial stiffening is greatest within one year of a woman’s final menstrual period, as compared with the 5 years preceding or following menopause^7^. Thus, progression through menopause, and ultimately the reduction in circulating estrogen levels^8^, likely contribute to female arterial stiffening with aging. However, the molecular mechanisms mediating this increase in cardiovascular risk remain unclear.

As estrogens primarily mediate the protective effects in the cardiovascular system by binding to estrogen receptors^9,10^, estrogen receptor-dependent signaling may impact arterial stiffening. There are three main receptors for estrogens, including estrogen receptor alpha (ERα), estrogen receptor beta (ERβ), and the G protein-coupled estrogen receptor (GPER)^11^. Prior investigations utilizing whole-body knockout mouse models identified that ERα specifically regulates the protective effects of estrogen against vascular injury^9,10^. Further, ERα has also been shown to directly contribute to vascular smooth muscle cell (SMC) function and arterial phenotypes^12,13^ relevant to arterial stiffening^14^. Additionally, there are human single nucleotide polymorphisms in the gene that encodes for ERα (ESR1) that are associated with protection from arterial stiffening in postmenopausal women compared with age-matched men^15,16^. These findings support both a clinical relevance of and mechanistic basis for a role of ERα specifically in the pathogenesis of arterial stiffening, potentially through SMC-specific functions, which is the primary cell type in arteries, including the aorta^17^. Further, there is evidence that ERα may elicit vascular protective effects independent of estradiol^12,16^, the prominent type of estrogen in premenopausal women^18,19^. Because estradiol levels are similar in postmenopausal women and age-matched men^8^, this positions ERα as a potential independent regulator of arterial stiffening in both women and men. However, the role of ERα in aging-associated arterial stiffening has not been investigated, especially the specific role of ERα in SMCs. Addressing these questions will help identify novel mechanisms of postmenopause- and aging-associated arterial stiffening, facilitating the development of new precision-based medicine strategies.

To address these important gaps in knowledge, we utilized an innovative SMC-specific ERα knockout mouse model to determine the role of estradiol and SMC-ERα in arterial stiffening under two conditions: 1) bilateral ovariectomy and 2) chronological aging. Our approach revealed a highly novel and sex-specific role of SMC-ERα in arterial stiffening in both females and males that is independent of estradiol.

## Methods

### Animal studies

All animals were handled in accordance with US National Institutes of Health standards, and all procedures were approved by the Tufts University Institutional Animal Care and Use Committee. Mice used for all studies were on a C57Bl6 background and were maintained in temperature- and humidity-controlled rooms, with 12 hr:12 hr light/dark cycles with ad libitum access to standard chow and water.

### Creation of novel SMC-specific ERα knockout mouse

We generated an inducible, SMC-specific, ERα knockout mouse colony on a C57Bl6 background by crossing an established floxed ERα mouse^9,20^ with SMA-Cre-ERT2 mice (smooth muscle actin promoter-driven expression of Cre-ERT2 recombinase that is activated by tamoxifen) as previously described^21,22^. All mice (male and female, Cre- and Cre+) were treated with 1 mg/kg/day of tamoxifen (Sigma) in sunflower oil (Sigma) for 5 days when mice were between 6 and 8 weeks of age to induce recombination of ERα in smooth muscle actin containing tissues.

### DNA PCR

At the time of euthanasia, tissues were frozen in liquid nitrogen. DNA was extracted using the DNeasy Tissue Kit (Qiagen), and polymerase chain reaction (PCR) was performed to examine recombination of genomic DNA from various tissues of SMA-Cre positive and negative animals post-tamoxifen induction, using primers (5’TTgCCCgATAACAATAACAT 3’ and 5’TgCAgCAgAAggTATTTgCCTgTTA3’) selected outside of loxP flanked DNA sequences to yield PCR products of 255 base pairs (floxed, recombined) or 815 base pairs (WT, not recombined).

### Experimental protocols

(A) Ovariectomy studies in wildtype mice. Female C57Bl6 mice were purchased from Jackson Laboratories (Bar Harbor, ME). At 8-10 weeks of age, mice were randomized to receive bilateral ovariectomy (OVEX) or sham (SHAM) operation. Two weeks later, a placebo or estradiol (E2, 0.36 mg/30 days)^23^ pellet (Innovative Research of America, Sarasota, FL) was implanted. This generated three randomized groups of mice (n=12-16/group): 1) SHAM + placebo pellet (SHAM+P), 2) OVEX + placebo pellet (OVEX+P), and 3) OVEX + E2 pellet (OVEX+E2). The final experimental endpoint occurred 6 weeks after the initial OVEX or SHAM surgery. (B) Ovariectomy studies in SMC-ERα knockout mice. At 10 weeks of age, female ERα intact (Cre-) and knockout (KO, Cre+) littermates were randomized to receive an OVEX or SHAM procedure (n=6-8/group), as described above. The final experimental endpoint occurred 6 weeks after the initial surgery. (C) Aging studies in SMC-ERα knockout mice. Female and male, SMC-ERα intact (Cre-) and KO (Cre+) littermates were studied at 3 and 18 months of age (n=9-13/group). These ages in mice model periods of hormonal availability in the human lifespan, such that female and male mice that are 3 months of age have achieved reproductive maturity, and that female mice are post-reproductive at 18 months of age^24^.

### Surgical procedures

For all surgical procedures, mice were anesthetized via inhalation of 3-5% and maintained at 1-3% isofluorane. (A) Ovariectomy/sham surgeries. Within each study, mice were randomized to receive either OVEX or SHAM operation, as previously described^25,26^. A 5 mm incision was made on the dorsal side, inferior to the lowest rib and parallel to the spine. The skin and muscle were dissected, and the fat pad and ovary were pulled through the incision. In the animals receiving the OVEX surgery, ovaries were then clamped, tied off with 6-0 silk ties, and each ovary was separated from the uterine horn with microsurgical scissors. (B) Subcutaneous pellet implantation. OVEX mice were randomized to the placement of a placebo or E2 pellet. Pellets were implanted subcutaneously via a 3 mm transverse incision on the dorsal side between the shoulders. A 10G trochar was used to insert the pellet, where it was then left in place for the remainder of the study. (C) Radiotelemeter implantation. A 1.5 cm midline incision was made on the ventral neck region, from the chin to the clavicle. The left common carotid artery was dissected, tagged with a 7-0 silk tie, and ligated at the distal end just proximal to the carotid bifurcation. The telemetric device was placed subcutaneously in the animal’s left flank. The catheter end of the device was placed into the carotid artery via a small transverse arteriotomy and the tip advanced to the level of the aortic arch (∼10 mm).

### Circulating estradiol measurements

Whole blood was kept on ice until centrifuged at 2.0 rcf for 20 min at 4°C. Serum was aliquoted and frozen at -80°C. Estradiol was quantified via liquid chromatography-mass spectrometry at the Immunochemical Core Laboratory at the Mayo Clinic in Rochester, MN.

### Aortic pulse wave velocity

*In vivo* aortic pulse wave velocity (PWV) was measured as an index of arterial stiffness using Doppler ultrasound (Vevo 2100, VisualSonics, Toronto, ON, Canada). Mice were anesthetized with isofluorane and placed in a recumbent position on a heated platform (37°C). Electrocardiography (ECG) recordings were made via paw contact with pad electrodes. Mice were maintained at 1-3% isofluorane during the procedure to maintain a heart rate of 400-450 beats/min. The transit time between the proximal and distal abdominal aorta was determined by averaging distances between the foot of the flow waveform and the R-wave of the ECG over at least ten cardiac cycles at each location. PWV (mm/ms) was calculated by dividing the distance (mm) by the difference in transit times (ms) obtained at each location, as previously described^21,27^.

### Aortic histology

Mouse thoracic aortas were fixed in 10% neutral buffered formalin, paraffin-embedded, and sectioned. Sections were stained with Masson’s trichrome, to quantify aortic fibrosis, and Verhoeff van Gieson, to quantify elastin degradation. For quantification of fibrosis, collagen content was estimated as percent fibrosis of the medial layer using computerized morphometric analysis software (Fiji software), as previously described^21,27^. For quantification of elastin degradation, sections were assessed for number of breaks, number of broken elastin layers, and fraction of the aortic circumference with elastin breaks and then scored according to those criteria (0-5), as previously published^28–30^. All histologic analyses were performed by blinded investigators.

### Arterial blood pressure

Arterial blood pressure was measured *in vivo* via radiotelemetry (Data Sciences International, TA1 1PA-C10) in subsets of wildtype mice in ovariectomy studies (n=3-5/group) and aged female and male SMC-ERα intact and KO mice (n=10-16/group). Blood pressure data was recorded over at least 48 hours and averaged over equal light and dark cycle durations.

### Atomic force microscopy

Freshly dispersed aortic SMCs were cultured for 5 days (passage 0) in DMEM medium supplemented with 10% FBS, 1% L-glutamine, and 1% sodium pyruvate at 37°C with 5% CO_2_ and 100% humidity. Aortic SMCs were then serum starved overnight, and atomic force microscopy (AFM) was performed on isolated aortic SMCs from young and aged male and female SMC-ERα-intact and KO mice. A MFP-3D-Bio AFM system (Asylum Research Inc.) was mounted on an Olympus IX-81 microscope and operated in contact mode to collect force curves at the cell surface. MLCT AFM probes (Bruker Nano Inc.) with spring constant range of 11∼16 pN/nm were functionalized with mouse fibronectin (0.1 mg/ml Innovative Research, Inc.) and interacted with the cell surface to measure aortic SMC cortical stiffness and integrin α5β1-fibronectin adhesion, as described previously^31^. For each experiment, eight cells were measured per mouse, with 50 force curves being performed on each cell. Force curves were subsequently processed using MatLab software to determine the cell stiffness and integrin α5β1-fibronectin adhesion probability, as previously described^31^.

### Immunoblotting

Frozen aortas were homogenized using lysis buffer (Roche). Total protein (30 µg) was mixed in sample buffer [62.5 mM Tris-HCl, pH 6.8, 2% SDS, 25% glycerol, 0.01% bromophenol blue] and separated on 4-15% gradient polyacrylamide gel (Bio-Rad), transferred to polyvinylidene difluoride membranes (Millipore), blocked with 5% milk in PBS-Tween or Intercept Blocking Buffer (LiCor) for 1 hour, and probed with the indicated primary antibody overnight at 4°C. This was followed by appropriate secondary antibody for 1 hour at room temperature and development with enhanced chemiluminescence reagent. Optical density values were assessed via Fiji software and expressed normalized to established housekeeping proteins, as indicated, and set as a fold change relative to a chosen control group. All antibodies are detailed in Table S1.

### Statistical analysis

Between group differences were assessed using unpaired T-test, Welch’s analysis of variance (ANOVA), one-way ANOVA, two-way ANOVA, or repeated-measures two-way ANOVA where appropriate, as indicated in each figure legend. Tukey post hoc testing, Bonferroni post hoc testing, or Dunnett’s T3 multiple comparisons test were used for post hoc pairwise comparisons where appropriate, as indicated in each figure legend. The level of significance was set at α=0.05 for all statistical tests. All data are presented as mean ± standard error of the mean (SEM).

## Results

### Ovariectomy-driven loss of estradiol promotes arterial stiffening and fibrosis

To explore the role of estradiol loss in arterial stiffening in a rodent model, young female mice underwent a SHAM or OVEX surgery, followed by randomization of OVEX mice to placebo (P) or E2 supplementation by subcutaneous pellet (Figure 1A). Efficacy of the experimental model was verified by the expected impact of estradiol on uterus mass (Figure 1B) and circulating E2 levels (Figure 1C), demonstrating a loss of estradiol with OVEX+P compared with SHAM+P and successful restoration of E2 via subcutaneous pellet in OVEX+E2 compared with OVEX+P mice. Animal body and organ masses are displayed in Table S2. To assess the time course of arterial stiffening after OVEX, arterial stiffness was measured in an initial subset of mice prior to surgery and 2-, 4-, and 6-weeks post-surgery. Arterial stiffness was significantly and consistently increased at 4-and 6-weeks post-surgery in OVEX+P mice, while no significant change in arterial stiffness was observed over time in SHAM+P or OVEX+E2 mice (Figure 1D). This experiment was then replicated, and OVEX+P mice consistently had greater arterial stiffening relative to both other groups (Figure 1E), such that OVEX induced arterial stiffening, but this was prevented with E2 supplementation. A similar pattern was observed when PWV was expressed as the change in arterial stiffness (ΔPWV) from before surgery to 6-weeks-post surgery for each individual mouse (Figure 1F). Arterial blood pressure was measured in a subset of mice that underwent the same experimental protocol, and there was no significant difference in the change in mean arterial pressure between 1-week- and 6-weeks post-surgery (calculated as the delta change for each individual mouse) between groups (Table S2), consistent with a direct effect of OVEX at the tissue level. Aortic sections revealed greater fibrosis in the medial layer of aortas from OVEX+P mice relative to SHAM+P mice, and this was completely prevented in OVEX+E2 mice (Figure 1G). There was no significant difference in aortic elastin degradation between groups (Figure 1H).

**Figure 1.**
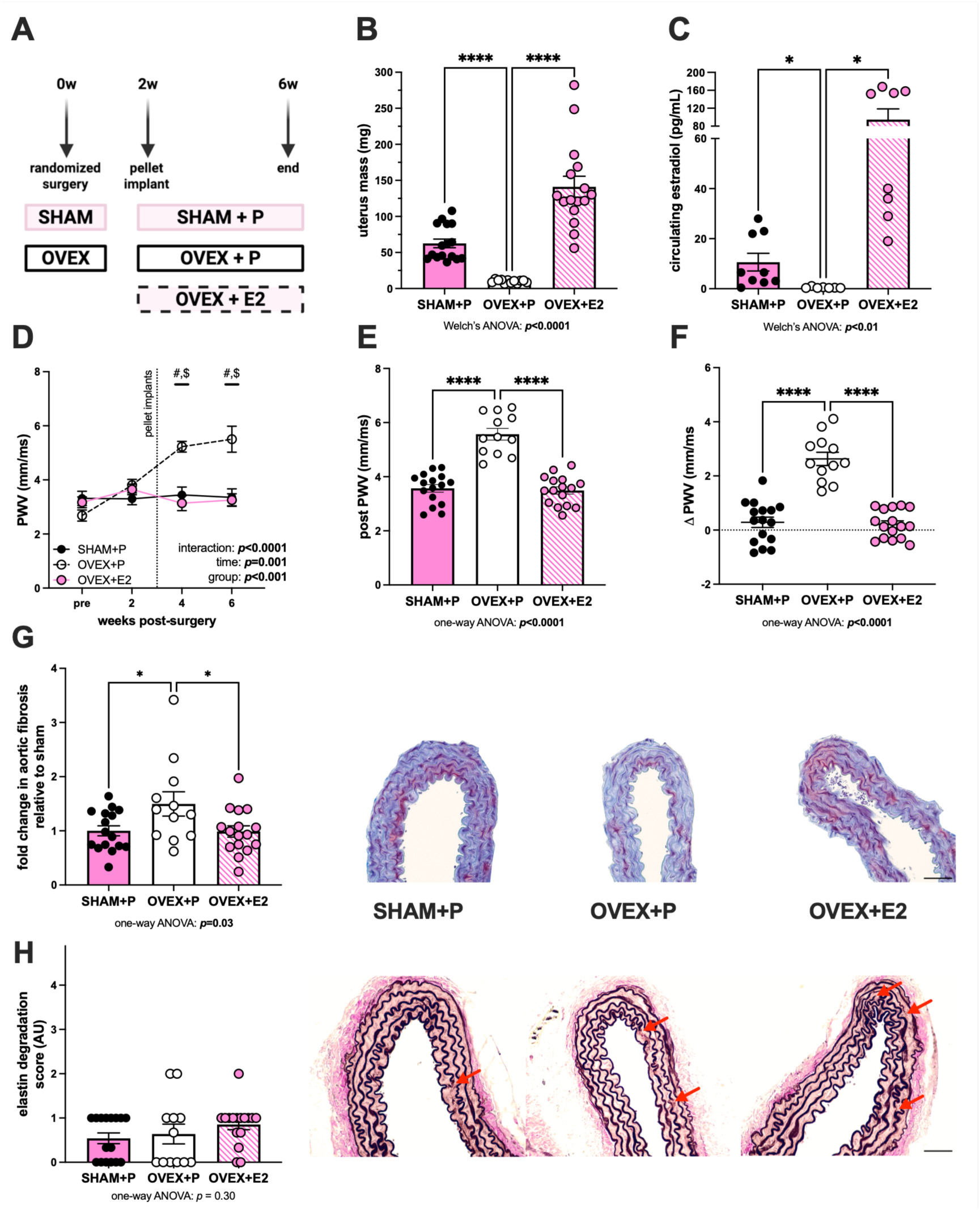
Ovariectomy-driven loss of estradiol promotes arterial stiffening and fibrosis. Young, female mice were randomized to three groups: sham surgery with placebo pellet (SHAM+P), bilateral ovariectomy with placebo pellet (OVEX+P), or OVEX with 0.36 mg estradiol pellet (OVEX+E2) (panel A). Successful OVEX surgery and E2 supplementation were confirmed by uterus mass (panel B) and circulating estradiol levels measured by liquid chromatography mass spectrometry (panel C). To determine a time course of arterial stiffening following OVEX, a subset (n=4-5/group) underwent serial pulse wave velocity (PWV) measurements before surgery and in 2-week increments following, until 6 weeks post-surgery (panel D). PWV was then replicated (n=12-16/group) and, group data is shown at 6 weeks post-surgery (panel E) and as delta values from individual mice (panel F). Histologic sections of aortas were stained with Masson’s trichrome to assess medial layer fibrosis (panel G) and stained with Verhoeff van Gieson to assess elastin degradation (panel H). Red arrows indicate elastin degradation. Scale bar is 10 μm. Group differences were assessed by Welch’s ANOVA (because of unequal variance, panels B and C), repeated measures two-way ANOVA (panel D), and one-way ANOVA (panels E-H). Dunnett’s T3 multiple comparisons (panels B and C) or Tukey (panels D-H) post-hoc testing were used. Data are means ± SEM, \**p*<0.05. \*\*\*\**p*<0.0001. #*p*<0.05 versus SHAM+P. $*p*<0.05 versus OVEX+E2.

### SMC-ERα contributes to ovariectomy-induced arterial stiffening and fibrosis

To investigate the role of SMC-ERα in arterial stiffening, we generated a novel mouse model with an inducible genetic KO of ERα from smooth muscle actin-containing tissues, including the aorta (our primary tissue of interest), bladder, and uterus, while the native ERα remained intact in tissues that are not predominantly composed of SMC (e.g., the liver, Figure 2A). To determine how the ligand status of SMC-ERα affects arterial stiffening, young female SMC-ERα-intact and KO littermates were randomized to receive a SHAM or OVEX surgery (Figure 2B). Successful OVEX surgery was confirmed by uterus mass, where uteruses from mice that underwent OVEX procedures were significantly smaller than those that had the SHAM procedure (Figure 2C). Animal body and organ masses are displayed in Table S3. There was greater arterial stiffening in mice that underwent OVEX procedure compared with those that received a SHAM procedure (Figure 2D), consistent with data in Figure 1 in wildtype mice. Deletion of ERα from SMC had no effect on arterial stiffening in mice that underwent SHAM surgery; however, the deletion of SMC-ERα prevented the OVEX-induced increase in arterial stiffening (Figure 2D), supporting that the presence of SMC-ERα in the absence of estradiol contributes to arterial stiffening. This pattern was reproducible when expressed as the ΔPWV from before surgery to 6-weeks-post surgery for each individual mouse (Figure 2E). In SMC-ERα-intact mice, OVEX promoted aortic fibrosis (Figure 2F), as in Figure 1G, but this was prevented in mice with the deletion of SMC-ERα (Figure 2G). The presence (Figure 2H) or absence (Figure 2I) of SMC-ERα had no impact on aortic elastin degradation following OVEX, recapitulating that OVEX does not significantly impact aortic elastin degradation within 6 weeks (Figure 1H).

**Figure 2.**
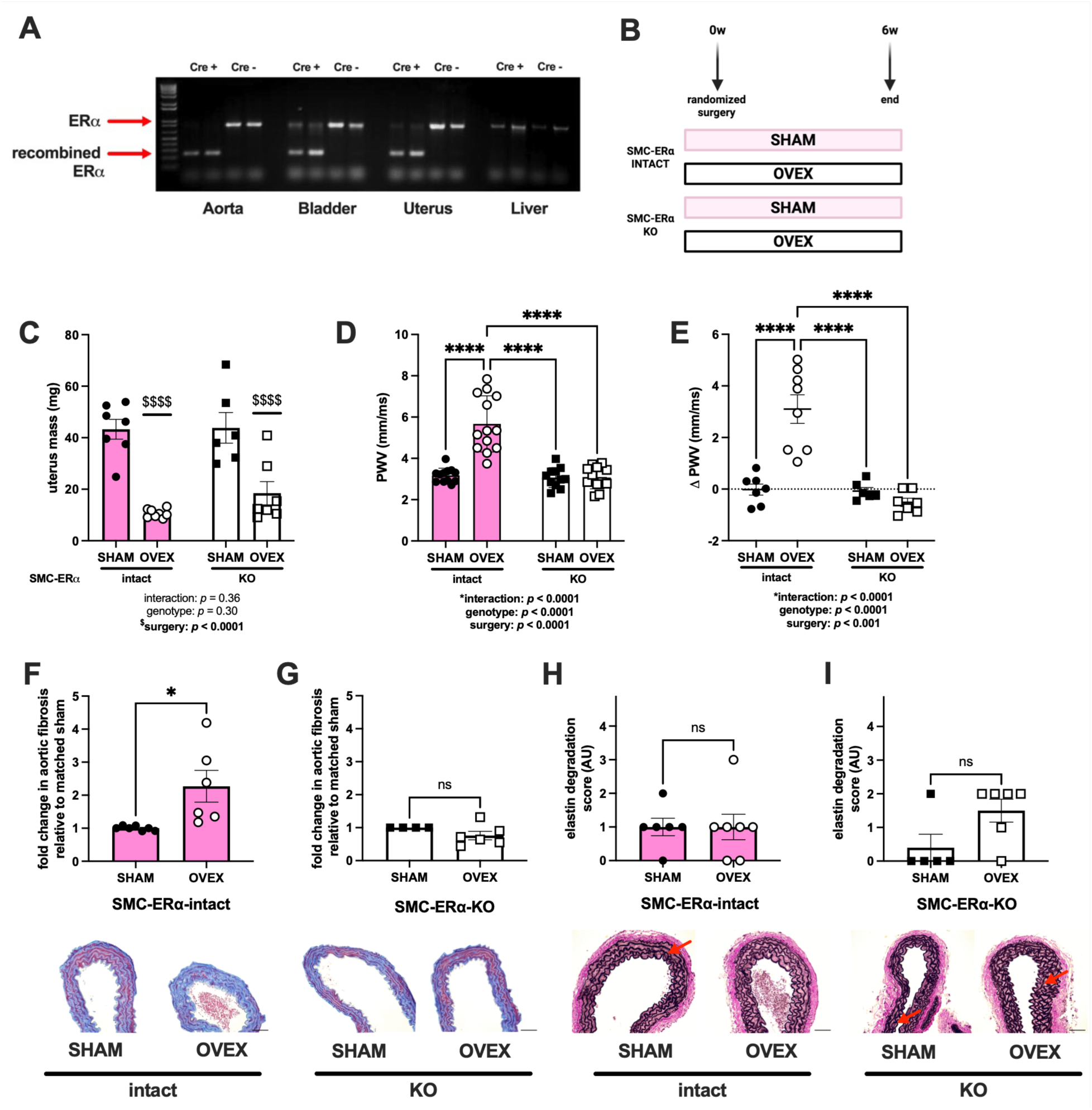
SMC-ERα contributes to ovariectomy-induced arterial stiffening and fibrosis. Smooth muscle cell (SMC)-specific estrogen receptor α (ERα) knockout (KO) mice were generated. SMC-ERα-intact (Cre-) and SMC-ERα-KO (Cre+) mice were generated. DNA PCR was used to show that Cre- mice displayed the native ERα in tissues containing and not containing SMC, while Cre+ mice displayed the presence of a recombined ERα in the tissues containing SMC but not tissues that are not primarily composed of SMC (i.e., the liver, panel A). Young, female SMC-ERα-intact and SMC-ERα-KO mice (n=6-8/group) were randomized to receive a sham (SHAM) or ovariectomy (OVEX) surgery and outcomes were measured after 6 weeks (panel B). Successful OVEX surgery was confirmed by uterus mass (panel C). Arterial stiffness was measured by pulse wave velocity (PWV) 6 weeks after surgical procedures (panel D). This pattern was replicated when PWV was assessed as a delta value from individual mice (panel E). Histologic sections of aortas were stained with Masson’s trichrome to assess medial layer fibrosis in SMC-ERα-intact (panel F) and SMC-ERα-KO (panel G) mice. Sections were stained with Verhoeff van Gieson to assess elastin degradation in SMC-ERα-intact (panel H) and SMC-ERα-KO (panel I) mice. Red arrows indicate elastin degradation. Scale bar is 10 μm. Group differences were assessed by two-way ANOVA (panels C-E) or unpaired T-test (panels F-I). Tukey’s post-hoc testing was used. Data are means ± SEM. \**p*<0.05. \*\*\*\**p*<0.0001. Main effect of age: $$$$*p*<0.0001. ns, not significant.

### SMC-ERα contributes to aging-associated arterial stiffening, but not fibrosis, in female and male mice

Chronological aging results in a decrease in circulating E2 in female mice (Figure 3A), and circulating E2 values consistently fall below the lower limit of detection in male mice^32,33^, supporting chronically low circulating E2 levels in male mice. To determine the role of SMC-ERα in arterial stiffening associated with chronological aging, and concurrent estrogen loss, we measured arterial stiffening in young and aged male and female SMC-ERα intact and KO mice (Figure 3B). Animal body and organ masses are shown in Table S4 (females) and S5 (males). Arterial stiffening was significantly greater in aged compared with young SMC-ERα-intact females (Figure 3C). There was no effect of the deletion of SMC-ERα on arterial stiffness in young females, replicating findings from Figure 2D; however, SMC-ERα-KO female mice were protected from the aging-associated increase in arterial stiffening (Figure 3C). Arterial stiffening was also significantly increased in aged compared with young SMC-ERα-intact males (Figure 3D). There was no effect of the deletion of SMC-ERα on arterial stiffness in young males, similar to that observed in young females. The aging-associated increase in arterial stiffening was partially prevented in SMC-ERα-KO males, such that arterial stiffness in aged SMC-ERα-KO males was significantly lower than that of aged SMC-ERα-intact males but remained greater than that of young male mice (Figure 3D). These findings further support that the presence of SMC-ERα in low estradiol environments promotes arterial stiffening.

**Figure 3.**
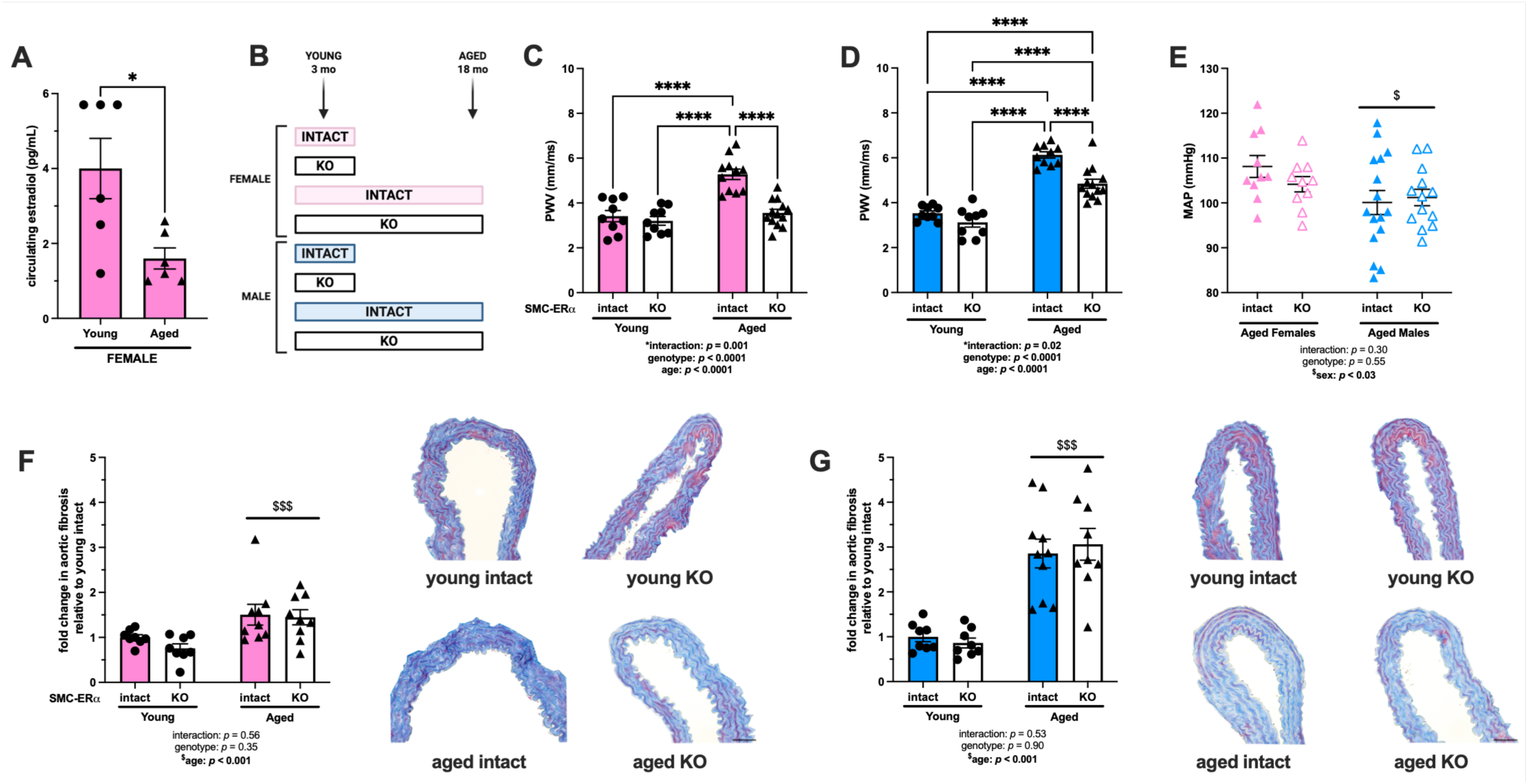
SMC-ERα contributes to aging-associated arterial stiffening, but not fibrosis, in female and male mice. Estradiol levels were measured by liquid chromatography mass spectrometry in young and aged female mice (panel A). Outcomes were measured in young and aged smooth muscle cell (SMC)-specific estrogen receptor α (ERα) intact and knockout (KO) female and male mice (panel B). Arterial stiffness was measured by pulse wave velocity (PWV) in female (panel C) and male (panel D) mice. *In vivo* 24-hr mean arterial pressure was measured by radiotelemetry in aged female and male SMC-ERα-intact and SMC-ERα-KO mice (panel E). Histologic sections of aortas were stained with Masson’s trichrome to assess medial layer fibrosis in female (panel F) and male (panel G) mice. Scale bar is 10 μm. Group differences were assessed by unpaired T-test (panel A) or two-way ANOVA (panels C-G). Tukey’s post-hoc testing was used. Data are means ± SEM. \**p*<0.05. \*\*\*\**p*<0.0001. Main effect of age: $*p*<0.05, $$$*p*<0.001.

Given the direct relationship between arterial stiffening and blood pressure, we next measured *in vivo* blood pressure via radiotelemetry in aged female and male SMC-ERα-intact and KO mice. In aged female and male mice, the presence or absence of SMC-ERα had no impact on mean arterial pressure (Figure 3E), again supporting a direct effect of SMC-ERα at the tissue level. However, aged males overall had significantly lower mean arterial pressure compared with aged females (Figure 3E) regardless of genotype. Aortic sections revealed an increase in aortic fibrosis in the medial layer of aortas from aged female (Figure 3H) and male (Figure 3I) mice compared with young counterparts, but there was no effect of the deletion of SMC-ERα, suggesting that ERα in SMCs regulates arterial stiffness induced by OVEX (Figures 1-2) and chronological aging by distinct mechanisms.

### SMC-ERα contributes to aortic elastin degradation in aged female, but not male, mice

Aortic sections revealed an increase in elastin degradation in aged compared with young female mice; however, there was an overall decrease in elastin degradation in female SMC-ERα-KO compared with SMC-ERα-intact mice (Figure 4A-B), consistent with the observation of reduced *in vivo* arterial stiffness in aged female SMC-ERα-KO mice (Figure 3C). Similar to females, there was an increase in elastin degradation in aged compared with young male mice; however, there was no effect of ERα deletion from SMC on this metric in male mice (Figure 4C, representative images Figure 4D).

**Figure 4.**
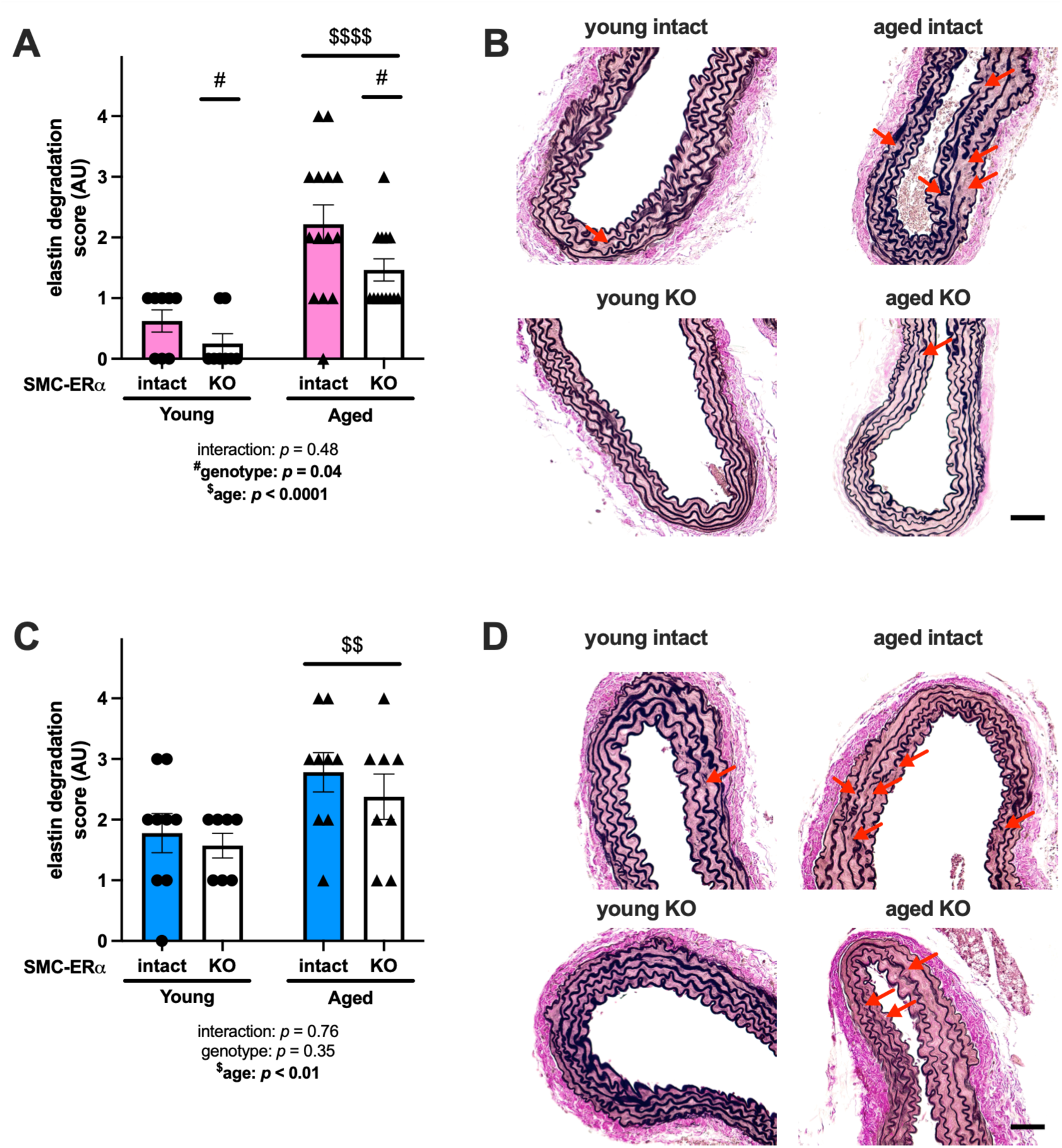
SMC-ERα contributes to aortic elastin degradation in female, but not male, mice. Outcomes were measured in young and aged smooth muscle cell (SMC)-specific estrogen receptor α (ERα) intact and knockout (KO) female and male mice. Histologic sections of aortas were stained with Verhoeff van Gieson to assess elastin degradation in female (panel A, representative images panel B) and male (panel C, representative images panel D) mice. Red arrows indicate elastin degradation. Scale bar is 10 μm. Group differences were assessed by two-way ANOVA (panels A and C). Bonferroni post-hoc testing was used. Data are means ± SEM. Main effect of genotype: #*p*<0.05. Main effect of age: $$*p*<0.01, $$$$*p*<0.0001.

### Sex-specific matrix metalloproteinase expression in aortas from SMC-ERα intact and knockout mice

Elastin degradation is mediated in part by matrix metalloproteinases (MMP); therefore, we next measured MMP protein expression in aortas from young and aged SMC-ERα-intact and SMC-ERα-KO mice. In female mice, MMP2 and MMP9 protein expression levels were increased in aortas from aged compared with young mice regardless of genotype (Figure 5A and 5B). However, MMP13 expression was greater in aortas from aged compared with young SMC-ERα-intact females, but there was no difference between MMP13 expression in aortas from aged and young SMC-ERα-KO females (Figure 5C). Further, MMP13 expression was greater in aortas from aged SMC-ERα-intact compared with aged SMC-ERα-KO female mice. This supports that MMP13 expression may contribute to SMC-ERα-mediated elastin degradation with aging in female mice. In aortas from male mice, MMP2 expression was increased in aortas from aged mice compared with young mice regardless of genotype (Figure 5D), and there were no statistical differences in aortic expression of MMP9 (Figure 5E) or MMP13 (Figure 5F) by age or genotype, correlating with a lack of change in elastin degradation phenotype when SMC-ERα is deleted in male mice.

**Figure 5.**
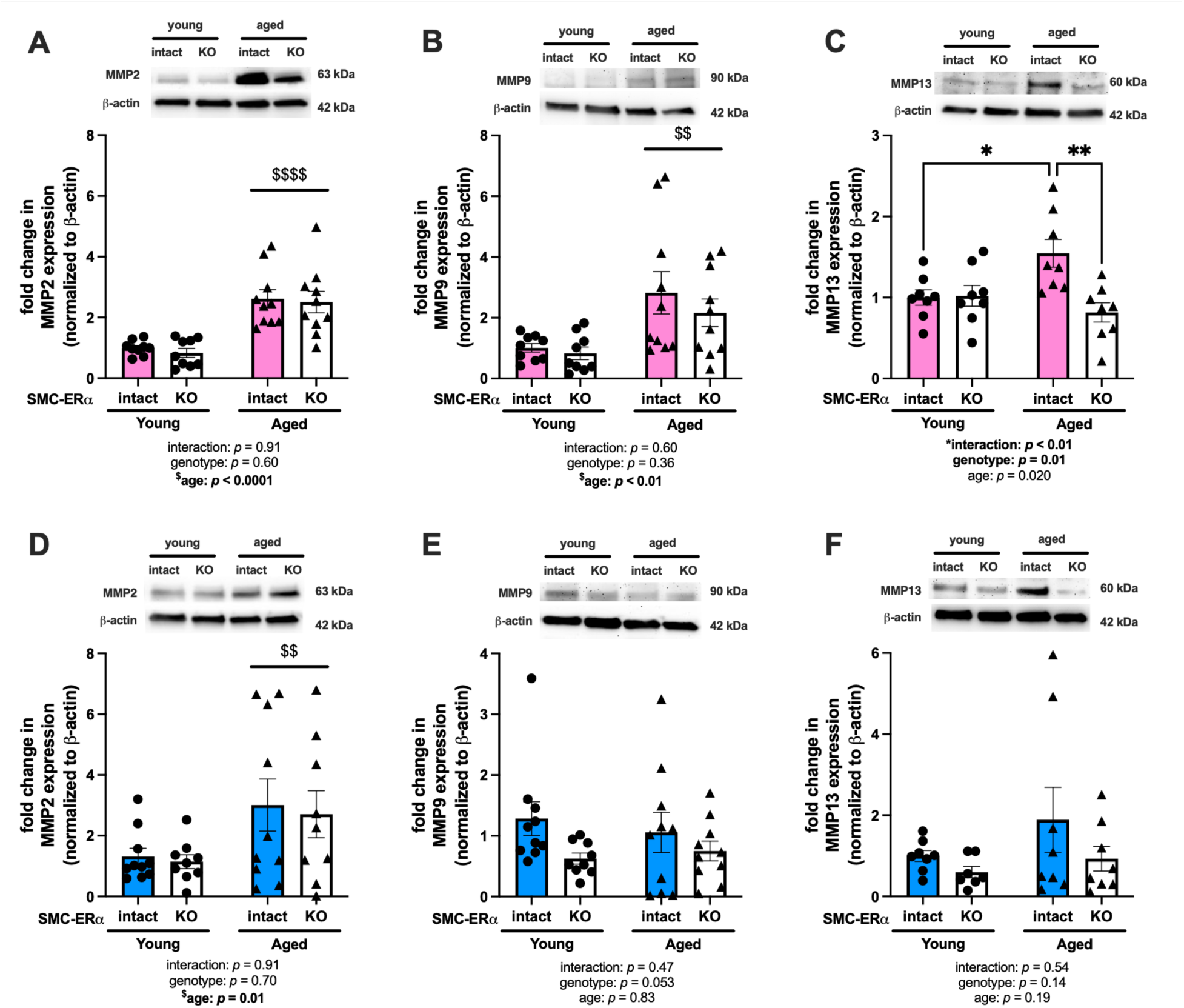
Sex-specific matrix metalloproteinase expression in aortas from SMC-ERα intact and knockout mice. Whole aortas were collected from young and aged smooth muscle cell (SMC)-specific estrogen receptor α (ERα) intact and knockout (KO) female and male mice and immunoblotted for protein expression. Expression of matrix metalloproteinase (MMP) 2 (panel A), MMP9 (panel B), and MMP13 (panel C) were measured in aortas from female mice and male mice (panels D-F, respectively). Group differences were assessed by two-way ANOVA (panels A-F). Bonferroni post-hoc testing was used. Data are means ± SEM. \*\**p*<0.01. Main effect of age: $$*p*<0.01, $$$$*p*<0.0001.

### ERα contributes to SMC stiffness and interactions with the extracellular matrix in aortic SMC from aged male, but not young or female, mice

To further investigate potential mechanisms by which SMC-ERα regulates aging-associated arterial stiffening, the intrinsic stiffness of aortic SMC and the probability of those aortic SMC to form adhesions with extracellular fibronectin were measured by AFM. There was no effect of the deletion of ERα from SMC on intrinsic stiffness of aortic SMC from young female (Figure 6A), aged female, (Figure 6B), or young male (Figure 6C) mice. Intrinsic stiffness was significantly reduced in aortic SMC from aged SMC-ERα-KO males compared with aged SMC-ERα-intact males (Figure 6D). There was no effect of the deletion of ERα from SMC on the probability of forming adhesions with fibronectin for aortic SMC from young female (Figure 6E), aged female (Figure 6F), or young male (Figure 6G) mice. In contrast, SMC from aged SMC-ERα-KO males were significantly less likely to form adhesions with fibronectin compared with SMC from aged SMC-ERα-intact males (Figure 6H). These findings are consistent with the observation of reduced *in vivo* arterial stiffness in aged male SMC-ERα-KO compared with SMC-ERα-intact mice (Figure 4D) and highlight an apparent sex-specific mechanism of action of SMC-ERα in aging-associated arterial stiffening.

**Figure 6.**
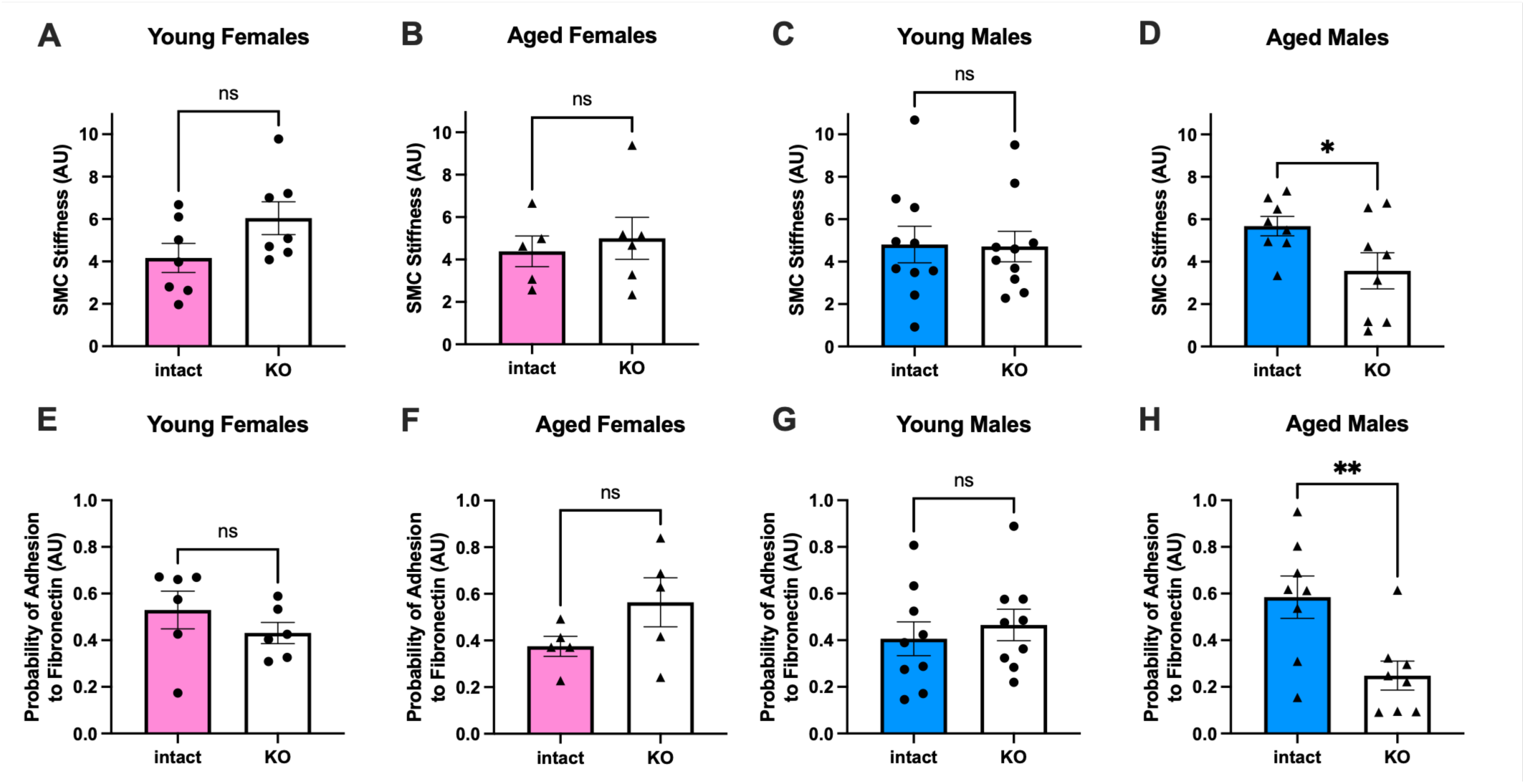
ERα contributes to SMC stiffness and interactions with the extracellular matrix in aortic SMC from aged male, but not young or female, mice. Intrinsic smooth muscle cell (SMC) stiffness (panels A-D) and the probability of forming adhesions with a fibronectin-coated probe (E-H) was measured by atomic force microscopy. Outcomes were measured in SMC from SMC-ERα-intact and SMC-ERα-KO young (panels A and E) and aged female mice (panels B and F), and young (panels C and G) and aged (panels D and H) male mice. Group differences were assessed by unpaired T-test (all panels). Data are means ± SEM. \**p*<0.05. ns, not significant.

Additional proteins associated with SMC stiffness were quantified by immunoblotting, including integrinβ1, integrinα5, focal adhesion kinase (FAK), and phosphorylation of FAK at tyrosine 397 (pFAK) in whole aortas from female and male SMC-ERα-intact and SMC-ERα-KO mice (Supplemental Figure S1). In aortas from female mice, integrinβ1 expression was lower in both SMC-ERα-KO mice and aged mice overall, with no interaction effect between factors. Additionally, integrinβ1 expression was greater in aortas from aged versus young SMC-ERα-KO male mice. Total FAK expression was also reduced in aortas from aged versus young female and male mice overall. There was no indication for changes in integrinα5 or pFAK expression between any groups of mice.

## Discussion

The major novel findings from the present study are as follows: (1) SMC-ERα promotes arterial stiffening in low estradiol environments, (2) SMC-ERα promotes arterial stiffening by distinct mechanisms in female mice that have undergone OVEX surgery versus chronological aging, and (3) SMC-ERα promotes aging-associated arterial stiffening in both female and male mice but does so by distinct, sex-specific mechanisms. Collectively, these findings support the novel concept that SMC-ERα significantly contributes to arterial stiffening in female and male mice in situations where the vasculature is exposed to low estradiol levels.

### SMC-ERα promotes arterial stiffening in low estradiol environments

Our findings are the first to show that SMC-ERα promotes arterial stiffening when circulating estradiol is low, such as in female mice after OVEX surgery and in chronologically aged female and male mice. There is established evidence in the clinical literature for an increase in arterial stiffening in women after menopause and significant impact of this on women’s cardiovascular risk and mortality thereafter^4,5,7,15,16^. Prior *in vitro* and preclinical literature has further provided evidence for estradiol-independent effects of ERα in SMC specifically and the vasculature in general^12,34^. Thus, SMC-ERα may provide a mechanistic link to vascular pathophysiology when circulating estradiol is low. Observations from *in vitro* experiments have revealed that when ERα in SMC is in the absence of estadiol, it contributes to alterations in gene expression consistent with vascular dysfunction, and that this gene expression profile is distinct from that observed in SMC when ERα is in the presence of estradiol^12^. In conjunction with the present data, it is thus plausible that ERα can influence gene expression, and therefore downstream protein translation, both in the presence and absence of estradiol (or other estrogens), which is a paradigm shift from classical understanding of the genomic functions of steroid hormones. Supporting this hypothesis, ERα has been shown to predominantly localize within the nucleus in various cell types regardless of the presence or absence of hormonal activation^35–38^, providing opportunity for ERα to contribute to physiology independent of estradiol. Additional hypotheses for the origin of estradiol-independent function of ERα include possible activation of ERα by alternate pathways, such as the phosphorylation of ERα via various growth factors^13,39,40^, thus generating ERα-dependent, estradiol-independent downstream effects. The present findings demonstrate evidence of SMC-ERα-dependent, estradiol-independent effects on *in vivo* vascular physiology, and future studies should focus on the precise mechanism(s) by which this occurs.

### SMC-ERα promotes arterial stiffening by distinct mechanisms in female mice following OVEX versus chronological aging

Our novel findings also show that SMC-ERα contributes to both OVEX- and aging-associated arterial stiffening in female mice, but that this appears to be induced by different mechanisms at the tissue level. We observed a reduction in aortic medial fibrosis accompanying the prevention of arterial stiffening in SMC-ERα-KO female mice following OVEX, but a reduction in elastin degradation accompanying the prevention of arterial stiffening in chronologically aged SMC-ERα-KO female mice. This context is important because neither surgical OVEX nor chronological aging in mice is precisely analogous to human menopause, though they do replicate certain aspects of the human condition and provide clues into the mechanisms mediating observations in our physiology. Recent preclinical studies have observed an increase in intracarotid arterial stiffness in both OVEX and 12-month-old female mice^41^; however, this study observed no change in aortic fibrosis following OVEX but increased aortic fibrosis in 12-month-old female mice. Similar to our data, this study further observed that neither OVEX- nor aging-associated arterial stiffening in female mice was directly related to increased arterial blood pressure^41^, further supporting that alterations in arterial stiffness are occurring at the tissue level.

This is the first study to thoroughly and directly investigate the loss of estradiol in arterial stiffening. Although the drastic reduction of circulating estradiol is a well-known outcome of the menopausal transition^8^, and there are clinical observations of an increase in arterial stiffening across the menopausal transition^7^, estradiol’s specific role in arterial stiffening after menopause has not been mechanistically interrogated in preclinical models. The menopausal transition is also accompanied by changes in circulating levels of several other hormones, including progesterone, follicle stimulating hormone, luteinizing hormone, and testosterone, among other changes. To date, five studies have investigated the effect of OVEX on arterial stiffening in rodent models^41–45^. These studies have yielded inconsistent results, indicating that OVEX increases^41,43^, decreases^44^, or has no effect^42,45^ on arterial stiffening. Our data here replicates recently published circulating E2 levels in young SHAM+P and OVEX+P female mice^41,42^. Here, we further causally link the loss of estradiol specifically with arterial stiffening in female mice after OVEX using an estrogen rescue experimental protocol and have completed a time course analysis to rigorously assess the temporal association of the loss of estradiol with arterial stiffening, further supporting that this is mediated by the presence and ligand status of SMC-ERα.

### SMC-ERα promotes aging-associated arterial stiffening in both female and male mice by sex-specific mechanisms

Here, we observe that SMC-ERα contributes to aging-associated arterial stiffening in female and male mice, but the mechanism by which this occurs is sex-specific. In 18-month-old SMC-ERα-KO compared with SMC-ERα-intact female mice, deletion of SMC-ERα reduced elastin degradation, and this was associated with a reduction in aortic MMP13 expression. MMP13 has previously been shown to be regulated by ERα in other cell types, especially in connective tissues^46–49^. Previous studies have also shown that ERα can enhance activity of the MMP13 promotor via binding at the AP-1 transcriptional regulatory site, and this effect is significantly depressed in the presence of estradiol^47^. Additionally, both the AF-1 and AF-2 domains of ERα appear to play a role in this mechanism, where ERα constructs lacking the AF-2 domain lead to significant elevation in MMP13 promoter activity, and those lacking the AF-1 domain lead to significant loss in MMP13 promoter activity^46^. Further, the binding of estrogens to ERα promotes a conformational change that leads to the exposure of the AF-2 domain, allowing complex formation with coactivators such as AP-1 and steroid receptor coactivator-1 (SRC-1)^50^. Collectively, this suggests that in the absence of estrogens, the AF-1 domain may be enhancing MMP13 promoter activity. Importantly, MMP13 is better known for its role as a collagenase than as an elastase^51^, though it does indirectly participate in elastin degradation through its role in inactivating alpha-1 antitrypsin (AAT), the body’s primary inhibitor of neutrophil elastase (NE)^52,53^. NE mRNA expression and activity in the aortic wall has been shown to be increased in aged compared with young mice^54^. Thus, SMC-ERα in the progressively induced low estradiol environment associated with chronological aging may increase MMP13 promoter activity and transcription, influencing elastin degradation and thus promoting arterial stiffening, consistent with the present observations in female mice within this study. Future studies are warranted to directly investigate the regulation of interaction between ERα and MMP13 in sex-specific situations and as a function of estradiol concentration. Additionally, AAT and NE may contribute to downstream mechanisms of arterial stiffness, and future studies may aim to dissect this potential mechanism in SMC, specifically.

In contrast to the observations in female mice in this study, the deletion of SMC-ERα had no influence on elastin degradation phenotype in male mice; however, the deletion of SMC-ERα reduced the intrinsic cortical stiffness and probability of extracellular matrix adhesion formation in aortic SMC from aged male mice, while deletion of SMC-ERα had no effect in aortic SMC from aged female mice. Intrinsic SMC stiffness and adhesion formation phenotypes may be influenced by proteins such as actin and myosin filaments, fibrillin-1, fibronectin, talin, integrins, FAK, paxillin, and cadherins^55^. While some studies have noted potential interactions of these proteins with ERα^56-59^, no investigations currently address these proteins specifically in vascular SMC. Previous observations support that the filamentous (F) actin to globular (G) actin ratio in vascular SMC contributes to hypertension-induced arterial stiffening^60,61^ and that ER subtypes can influence actin polymerization in pulmonary SMC^56^ and vascular endothelial cells^57^. Additionally, ERα regulates phosphorylation and complex formation of FAK in human umbilical vein endothelial cells, ultimately promoting motility of endothelial cells^58^. Integrinβ1 and fibronectin have been shown to co-localize with ERα in human breast cancer cells, and the presence of fibronectin reduces the lysosomal degradation of both integrinβ1 and ERα in the presence of E2^59^. The protein expression data presented here (Figure S1) provides initial indication for further investigation. Future work should focus on mechanistic links between SMC-ERα and stiffness-associated proteins to understand sex- and aging-dependent contributions to arterial stiffening.

### Limitations

Neither the surgical OVEX nor chronological aging models in mice recapitulate all of the physiology associated with human menopause, but there are similarities that support the importance of these findings. These models allowed for the dissection of the specific effects of the loss of estradiol when paired with surgical pellet implantation and of ERα in SMC on OVEX- and aging-induced arterial stiffening. Importantly, mice do not undergo an equivalent process to human menopause, but new experimental models, such as the 4-vinylcyclohexene diepoxide (VCD) model, may better replicate certain aspects of human menopause that are not captured in the models used in this study. As such, future studies may implement the VCD design, which highlights a focus on follicle atresia within intact ovaries, which may impact circulating levels of other hormones, for further exploration of the mechanisms underlying menopause-induced arterial stiffening.

## Conclusions

Overall, this study provides evidence to support that ERα in SMC contributes to arterial stiffening in female and male mice when the vasculature is exposed to low estradiol concentrations. This occurs mechanistically through sex-specific pathways in response to chronological aging and is mediated by different pathways in response to a reduction in estradiol in females due to OVEX surgery versus chronological aging. These findings implicate a role of lower circulating levels of estradiol and the ligand status of ERα in major cardiovascular complications in women and men observed with aging and/or menopause. Future studies should further interrogate the mechanism of action of SMC-ERα on the development of these outcomes to influence precision-based medicine and contribute to the identification of novel therapeutic targets for CVD.

## Abbreviations

AAT: alpha-1 antitrypsin
AFM: atomic force microscopy
E2: estradiol
ECG: electrocardiogram
ERα: estrogen receptor alpha
ERβ: estrogen receptor beta
GPER: G protein-coupled estrogen receptor
KO: knockout
MMP: matrix metalloproteinase
NE: neutrophil elastase
OVEX: ovariectomy surgery
PWV: pulse wave velociy
SHAM: sham surgery
SMC: smooth muscle cell

## Acknowledgments

The authors thank Nathan Li and Terance Cheung with the Tufts Animal Histology Core for their technical help with aortic histology.

## Author Contributions

CGT and JJD conceived the research. CGT, JM, JB, KCO, RK, JV, MZ, JI, QL, GM, and ZS performed experiments. CGT, JM, JB, and ZS analyzed data. CGT, JM, JB, QL, ZS, IZJ, MAH, and JJD interpreted data. CGT, JM, and MZ prepared figures. CGT drafted the manuscript. CGT, JM, JB, KCO, RK, JV, MZ, JI, QL, GM, ZS, IZJ, MAH, and JJD revised the manuscript and approved the final version of this manuscript.

## Supplemental Materials

Figure S1.

Table S1-S5.

## HIGHLIGHTS

- Estrogen receptor alpha (ERα) in smooth muscle cells (SMC) promotes arterial stiffening in low estradiol environments.
- SMC-ERα promotes arterial stiffening by distinct mechanisms in female mice that have undergone a reduction of circulating estradiol via an ovariectomy surgery versus chronological aging.
- SMC-ERα promotes aging-associated arterial stiffening in both female and male mice through sex-specific mechanisms.

