## Supplemental Material for "Smooth muscle cell estrogen receptor alpha promotes arterial stiffness in the absence of estradiol"

**Affiliations**
[1]Molecular Cardiology Research Institute, Tufts Medical Center, Boston, MA, USA
[2]School of Kinesiology, Auburn University, Auburn, AL, USA
[3]College of Allopathic Medicine, Nova Southeastern University, Fort Lauderdale, FL, USA
[4]Translational Cardiology Unit, Navarrabiomed-Hospital Universitario de Navarra (HUN)-Universidad Pública de Navarra (UPNA), IDISNA, Pamplona, Navarre, Spain
[5]Dalton Cardiovascular Research Center, University of Missouri, Columbia, MO, USA

**Short Title:** Role of SMC-ER$\alpha$ in arterial stiffening

***Corresponding Author:**

Jennifer J. DuPont

Tufts Medical Center

800 Washington Street, Box 80

Boston, MA 02111

617 636-0620

**Keywords:** arterial stiffness, aging, estradiol, estrogen receptor alpha, sex differences

### SUPPLEMENTAL FIGURES

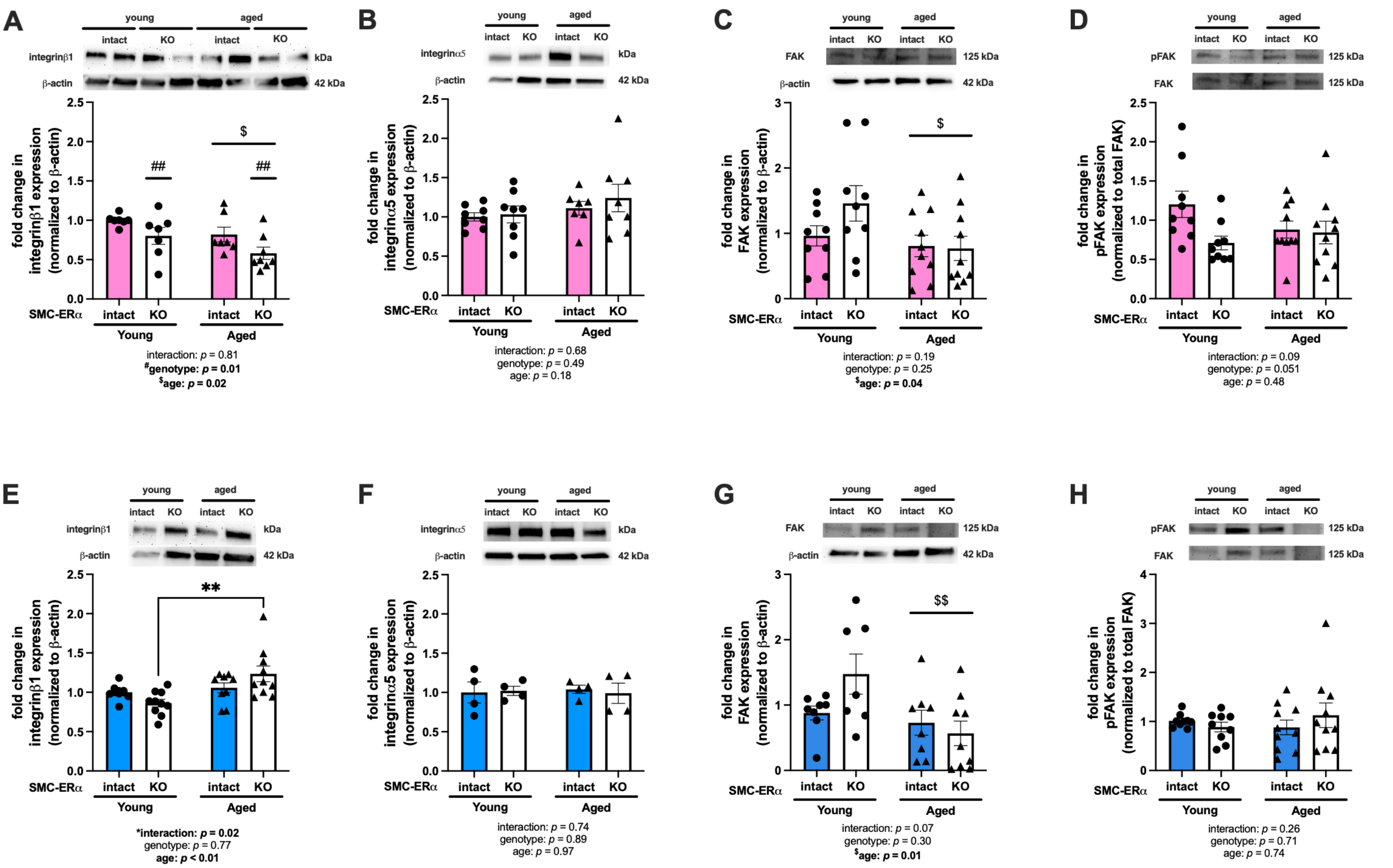


**Figure S1. Sex-specific expression of SMC-stiffness associated proteins in aortas from SMC-ERα intact and knockout mice.** Whole aortas were collected from young and aged smooth muscle cell (SMC)-specific estrogen receptor α intact and knockout (KO) female and male mice and immunoblotted for expression of proteins associated with SMC stiffness. Expression of integrinβ1 (panel A), integrinα5 (panel B), focal adhesion kinase (FAK, panel C), and FAK phosphorylated at tyrosine 397 (pFAK, panel D) were measured in aortas from female mice and aortas from male mice (panels E-H, respectively). Group differences were assessed by two-way ANOVA (panels A-H). Bonferroni post-hoc testing was used. Data are means ± SEM. ***p*<0.01. Main effect of genotype: ##*p*<0.01. Main effect of age: $*p*<0.05, $$*p*<0.01.

#### SUPPLEMENTAL TABLES

**Table S1. Antibodies used for immunoblotting.**

| Target antigen | Vendor or Source | Catalog # |
|---|---|---|
| Anti-mouse | Cell Signaling | 7076 |
| Anti-rabbit | Cell Signaling | 7074 |
| β-actin | Proteintech | 20536-1-AP |
| FAK | Cell Signaling | 3285 |
| Integrinα5 | Abcam | ab226816 |
| Integrinβ1 | Abcam | ab179471 |
| MMP2 | Santa Cruz | sc-13594 |
| MMP9 | Santa Cruz | sc-21733 |
| MMP13 | Abcam | ab39012 |
| pFAK | Cell Signaling | 3283 |

**Table S2. Animal characteristics — Wildtype ovariectomy experiment.**

| Variable | SHAM + P | OVEX + P | OVEX + E2 |
|---|---|---|---|
| Total n | 16 | 12 | 16 |
| n with tibia length and kidney mass | 9 | 6 | 10 |
| n with blood pressure measurements | 5 | 3 | 5 |
| Δ 24-hr mean arterial pressure (mmHg) (6wk post-surgery minus 1wk post-surgery) | 3.0 ± 3.4 | 2.2 ± 1.7 | 0.1 ± 1.5 |
| Body mass 6wk post-surgery (g) | 22.0 ± 0.4$^{\#}$ | 26.3 ± 0.5$^{*,\#}$ | 24.0 ± 0.3 |
| Δ Body mass (g) | 2.7 ± 0.3$^{\#}$ | 6.8 ± 0.4$^{*,\#}$ | 5.1 ± 0.4 |
| Heart mass (mg) | 119 ± 3 | 125 ± 3 | 115 ± 1 |
| Left ventricular mass (mg) | 82 ± 2 | 85 ± 2 | 80 ± 1 |
| Tibia length (mm) | 17.3 ± 0.1 | 17.9 ± 0.2 | 17.6 ± 0.1 |
| Heart mass (mg) / tibia length (mm) | 7.1 ± 0.3 | 6.8 ± 0.2 | 6.6 ± 0.1 |
| Left ventricle mass (mg) / tibia length (mm) | 4.9 ± 0.2 | 4.7 ± 0.2 | 4.5 ± 0.1 |
| Average kidney mass (mg) | 140 ± 4 | 134 ± 6 | 148 ± 4 |
| Average kidney mass (mg) / tibia length (mm) | 8.1 ± 0.3 | 7.5 ± 0.4 | 8.4 ± 0.2 |

E2, estradiol pellet. g, gram. mg, milligram. mm, millimeter. P, placebo pellet. OVEX, ovariectomy surgery. SHAM, sham surgery. wk, week. *$p$<0.05 versus SHAM+P. $^{\#}p$<0.05, versus OVEX+E2.

**Table S3. Animal characteristics — SMC-ERα ovariectomy experiment.**

| **Variable** | **SHAM SMC-ERα-intact** | **OVEX SMC-ERα-intact** | **SHAM SMC-ERα-KO** | **OVEX SMC-ERα-KO** |
|---|---|---|---|---|
| Total n | 7 | 8 | 6 | 7 |
| Body mass 6wk post-surgery (g) | 22.2 ± 0.5 | 23.6 ± 0.7* | 21.6 ± 0.6 | 25.4 ± 1.2* |
| Δ Body mass (g) | 2.3 ± .5 | 4.1 ± .4* | 2.6 ± $0.6^{\#}$ | 5.3 ± $0.9^{*,\#}$ |
| Tibia length (mm) | 17.9 ± 0.1 | 18.0 ± 0.1 | 17.6 ± 0.2 | 17.9 ± 0.1 |
| Heart mass (mg) | 122 ± 5 | 116 ± 5 | 119 ± 7 | 125 ± 5 |
| Heart mass (mg) / tibia length (mm) | 6.8 ± 0.3 | 6.5 ± 0.3 | 6.8 ± 0.3 | 7.0 ± 0.2 |
| Left ventricular mass (mg) | 83 ± 3 | 80 ± 3 | 81 ± 5 | 85 ± 3 |
| Left ventricle mass (mg) / tibia length (mm) | 4.7 ± 0.1 | 4.4 ± 0.2 | 4.6 ± 0.3 | 4.8 ± 0.2 |
| Average kidney mass (mg) | 137 ± 4 | 125 ± 4 | 125 ± 6 | 131 ± 6 |
| Average kidney mass (mg) / tibia length (mm) | 7.7 ± 0.2 | 7.0 ± 0.3 | 7.1 ± 0.3 | 7.3 ± 0.3 |

ERα, estrogen receptor alpha. g, gram. KO, knockout. mg, milligram. mm, millimeter. OVEX, ovariectomy surgery. SHAM, sham surgery. SMC, smooth muscle cell. *$p<0.05$, main effect of surgery. $^{\#}p<0.05$, main effect of genotype.

**Table S4. Animal characteristics — Aging SMC-ERα females.**

| Variable | YOUNG SMC-ERα-intact | YOUNG SMC-ERα-KO | AGED SMC-ERα-intact | AGED SMC-ERα-KO |
|---|---|---|---|---|
| Total n | 9 | 9 | 17 | 11 |
| Body mass (g) | 22.7 ± 0.5 | $21.2 \pm 0.6^{\#}$ | 32.1 ± 1.2* | $28.5 \pm 1.6^{*,\#}$ |
| Tibia length (mm) | 17.6 ± 0.1 | 17.4 ± 0.1 | 17.9 ± 0.1* | 17.7 ± 0.1* |
| Heart mass (mg) | 117 ± 7 | 111 ± 6 | 137 ± 6* | 138 ± 5* |
| Heart mass (mg) / tibia length (mm) | 6.7 ± 0.4 | 6.4 ± 0.3 | 7.6 ± 0.3* | 7.8 ± 0.3* |
| Left ventricular mass (mg) | 80 ± 5 | 78 ± 4 | 92 ± 3* | 94 ± 4* |
| Left ventricle mass (mg) / tibia length (mm) | 4.5 ± 0.2 | 4.5 ± 0.2 | 5.1 ± 0.2* | 5.3 ± 0.2* |
| Average kidney mass (mg) | 119 ± 6 | 118 ± 4 | 163 ± 7* | 163 ± 7* |
| Average kidney mass (mg) / tibia length (mm) | 6.8 ± 0.3 | 6.7 ± 0.2 | 9.1 ± 0.4* | 9.2 ± 0.4* |
| Uterus mass (mg) | 40 ± 7 | 34 ± 4 | 113 ± 14* | 73 ± 17* |
| Uterus mass (mg) / tibia length (mm) | 2.3 ± 0.4 | 1.9 ± 0.2 | 6.3 ± 0.8* | 4.1 ± 0.1* |

ERα, estrogen receptor alpha. g, gram. KO, knockout. mg, milligram. mm, millimeter. SMC, smooth muscle cell. *$p<0.05$, main effect of age. $^{\#}p<0.05$, main effect of genotype.

**Table S5. Animal characteristics — Aging SMC-ERα males.**

| Variable | YOUNG SMC-ERα-intact | YOUNG SMC-ERα-KO | AGED SMC-ERα-intact | AGED SMC-ERα-KO |
|---|---|---|---|---|
| Total n | 10 | 9 | 10 | 8 |
| Body mass (g) | 31.4 ± 1.2 | 28.3 ± 0.7 | 41.4 ± 2.6* | 39.2 ± 1.8* |
| Tibia length (mm) | 18.1 ± 0.1 | 18.1 ± 0.1 | 18.3 ± 0.1* | 18.4 ± 0.2* |
| Heart mass (mg) | 158 ± 7 | 146 ± 6 | 178 ± 11* | 160 ± 4* |
| Heart mass (mg) / tibia length (mm) | 8.7 ± 0.4 | 8.0 ± 0.4 | 9.7 ± 0.6 | 8.7 ± 0.3 |
| Left ventricular mass (mg) | 104 ± 4 | 96 ± 4 | 122 ± 8* | 108 ± 5* |
| Left ventricle mass (mg) / tibia length (mm) | 5.8 ± 0.3 | 5.3 ± 0.2 | 6.7 ± 0.4* | 5.9 ± 0.2* |
| Average kidney mass (mg) | 161 ± 7 | 150 ± 5$^{\#}$ | 210 ± 8* | 182 ± 5$^{*,\#}$ |
| Average kidney mass (mg) / tibia length (mm) | 8.9 ± 0.4 | 8.3 ± 0.3$^{\#}$ | 11.5 ± 0.4* | 9.9 ± 0.3$^{*,\#}$ |

ERα, estrogen receptor alpha. g, gram. KO, knockout. mg, milligram. mm, millimeter. SMC, smooth muscle cell. *$p$<0.05, main effect of age. $^{\#}p$<0.05, main effect of genotype.
